# PFLISA: A New Ultrasensitive, Multiplex Immunoassay Platform Enables Cytokine Network Profiling in Sepsis

**DOI:** 10.64898/2026.11.23.768466

**Authors:** Minsu Jang, Anushree Seth, Scott L. Crick, Ubaid Ullah, Delia F. Tifrea, Marlon Sy, Abraham J. Qavi

## Abstract

Immune responses emerge from coordinated cytokine networks, but quantitative profiling remains constrained by limited clinical sample volumes and the very low circulating concentrations of many biologically important mediators. We introduce the Plasmonic Fluor-Linked ImmunoSorbent Assay (PFLISA), that combines plasmonic-enhanced fluorescence with spatially encoded antibody arrays in a standard 96-well format. PFLISA quantified 12 inflammatory cytokines in parallel from 15 μL of plasma, with lower limits of quantification of 3.4–110 fg/mL and analyte-specific quantitative ranges spanning multiple orders of magnitude. Within-well replicate spots showed an average coefficient of variation of 5.7%, and 35 of 36 analyte-by-quality-control conditions exhibited intra- and interplate coefficients of variation below 20% across 16 plates. We applied the platform to plasma from 100 patients with sepsis and 97 healthy controls as a clinically demanding demonstration. Eleven of the 12 cytokines were significantly elevated in sepsis after false-discovery-rate correction. Pro- and anti-inflammatory composite scores increased 10.6-fold and 3.9-fold, respectively, yielding a 2.0-fold increase in their ratio and revealing concurrent but disproportionate activation of the two responses. Gram-negative infections were associated with higher IL-6, IL-8, TNF-α, and IL-10 concentrations than Gram-positive infections. PFLISA thus brings low-abundance regulatory cytokines into the same patient-level profile as more abundant inflammatory mediators, increasing the portion of the immune network that can be quantitatively observed from limited clinical samples.

## INTRODUCTION

Immune responses are coordinated through networks of soluble signals rather than isolated biomarkers. Cytokines regulate immune-cell activation, recruitment, differentiation, and resolution, and the relative balance among these mediators can shape disease trajectory and clinical outcome (*1*). Dysregulation of these networks is implicated in sepsis (*2, 3, 4*), cancer (*5, 6*), autoimmune disorders (*7*), and infectious diseases (*8, 9, 10*). A fundamental challenge in measuring these systems is that their components can exist at concentrations separated by several orders of magnitude. Highly abundant inflammatory mediators may coexist with regulatory cytokines present at far lower concentration, even within the same sample (*11*). Analytical sensitivity therefore determines which portions of the immune network are quantitatively observable. When low-abundance mediators fall below the measurable range of an assay, the resulting profile becomes weighted towards the most abundant signals and provides only a partial representation of the underlying immune response. Network-level profiling consequently requires not only multiplexing, but sufficient sensitivity and dynamic range to measure low- and high-abundance immune mediators from the same specimen.

PFLISA™ (Plasmonic Fluor-Linked ImmunoSorbent Assay) is a novel assay platform from Brightest Bio that combines plasmonic-enhanced fluorescence signal amplification and spatially encoded antibody arrays to achieve ultrasensitivity, multiplexing and high dynamic range providing a means to expand the portion of the immune network that can be quantitatively observed from limited clinical samples (*12, 13*). In PFLISA, engineered metallic nanostructures called Plasmonic Fluors™ amplify excitation and modify fluorophore emission while retaining the specificity and workflow of conventional sandwich immunoassays (*13, 14, 15, 16*). Herein we used the PFLISA-Plex Human Inflammation Panel A that allows quantification of 12 cytokines in parallel within a single well of a 96-well plate down to single digit fg/mL levels (*17*). Ultrasensitivity, wide dynamic range, and multiplexing serve complementary roles in interrogating cytokine networks in health and disease. Ultrasensitivity allows measurement of low-abundance mediators. Broad dynamic range allows quantitation of these mediators as they can differ in concentration by orders of magnitude in health versus disease. Multiplexing allows interrogation of multiple mediators of the immune response simultaneously. Quantified across a 4-5 log dynamic range from 15 µL of sample, these capabilities increase the network coverage achievable from a single limited specimen (*13, 17, 18*).

These capabilities position the PFLISA assay not simply as a more sensitive cytokine assay, but as a tool for expanding the measurable immune network and resolving coordinated immune states. Sepsis provides a demanding test case because it is a system-level disorder in which multiple inflammatory and regulatory pathways are activated simultaneously across wide, patient-specific concentration ranges. Sepsis is a life-threatening syndrome caused by a dysregulated host response to infection and can progress to multi-organ dysfunction (*2*). Its diverse infectious sources and heterogeneous clinical presentation make single-analyte measurements inherently incomplete. This heterogeneity is expressed not only in which cytokines are elevated, but also in their relative magnitude and coordination across patients. Sepsis is now recognized less as a single disorder than as a heterogeneous syndrome encompassing distinct host-response phenotypes. This heterogeneity is widely invoked to explain the repeated failure of immunomodulatory trials, in which patients with opposing immune states have been enrolled under a single syndromic definition and treated as one population; therapies matched to the underlying response will therefore require patient-level immune phenotyping (*19, 20, 21*). It is critical then to better understand biomarker signatures in septic patients to allow discrimination of these patients into appropriate phenotypes. A platform that measures cytokines over a broad concentration range together can instead reveal the balance, coordination, and heterogeneity of the underlying immune response, rather than only its most abundant signals.

Sepsis is characterized by dysregulated and frequently concurrent activation of pro- and anti-inflammatory signaling pathways. Pro-inflammatory cytokines such as IL-1β, IL-6, and TNF-α promote leukocyte recruitment, endothelial activation, and antimicrobial defense (*2, 22*), whereas mediators including IL-4, IL-10, and IL-13 can constrain excessive inflammation. Rather than occurring as strictly sequential phases, these responses may coexist, and their relative magnitude and coordination may provide a more complete representation of immune state than either response alone (*23, 24*). Resolving this balance requires quantitative access to both highly elevated inflammatory mediators and lower-abundance regulatory signals within the same sample. This complex immune dysregulation is associated with multi-organ failure and substantial mortality in sepsis (*22, 25*).

Because immune states vary across patients and over the course of illness, attempts to modulate individual cytokines have produced inconsistent clinical results (*4, 26*). This limitation creates a measurement need: immune status must be represented as a coordinated multianalyte pattern rather than inferred from one marker. Capturing that profile requires quantitative access to both dominant inflammatory signals and lower-abundance regulatory mediators. A platform capable of quantifying multiple pro- and anti-inflammatory signals in parallel could support more complete patient stratification and, with serial sampling, longitudinal immune monitoring.

Numerous methods for cytokine measurement have been developed, but each captures a different fraction of the immune response. Enzyme-linked immunosorbent assays (ELISAs) offer high sensitivity and specificity but typically measure one analyte per well (*27*), requiring limited samples to be divided across multiple assays for network-level profiling. Bead-based platforms such as Luminex and cytometric bead arrays increase multiplexing (*28, 29*), but assay-dependent sensitivity can limit measurement of low-abundance cytokines. Single-molecule array (Simoa) methods provide exceptional sensitivity, demonstrating the value of extending cytokine measurements into the sub-pg/mL range, but multiplexing breadth, sample requirements, and detectability remain assay dependent. Newer massively multiplexed systems such as NULISA and Olink provide very high plexing, high sensitivity, broad dynamic range, and low sample volume requirements, but have traditionally suffered from reproducible quantitation, and are so costly from an instrumentation and consumables perspective as to be out of reach for most researchers. The unmet analytical need is an integrated platform that combines ultrasensitivity with multiplexing, has a high dynamic range, is in a format compatible with laboratory automation, and is affordable to individual researchers.

We present the PFLISA 12-plex Human Inflammation Panel A, an ultrasensitive multiplexed immunoassay enabling simultaneous quantification of twelve key inflammatory cytokines: IL-1α, IL-1β, IL-2, IL-4, IL-5, IL-6, IL-8, IL-10, IL-13, IFN-γ, IL-12p70, and TNF-α. Leveraging Brightest Bio’s Plasmonic Fluor technology, PFLISA combines fg/mL sensitivity with a broad dynamic range spanning 4-6 orders of magnitude, allowing low-abundance cytokines to be measured alongside highly elevated inflammatory mediators within the same assay (*13*). The assay requires 15 μL of plasma and measures all 12 cytokines within a single well of a 96-well plate.

Here, we tested whether low-volume, ultrasensitive multiplexing could translate into broader quantitative coverage of the immune response. We applied this assay to plasma from 100 patients with sepsis and 97 healthy controls. Rather than evaluating each cytokine only as an isolated biomarker, we examined composite inflammatory balance, patient-level heterogeneity, pathogen-associated profiles, and correlations among cytokines. These analyses allowed us to ask whether extending sensitivity across the cytokine concentration spectrum could reveal relationships among immune mediators that would be incompletely represented by the most abundant signals alone. This study therefore evaluates the central value of the platform: recovering multiple interacting components of an immune network from limited sample volume within one scalable assay.

## RESULTS

### Low-Volume, Single-Well Multiplexing and Analytical Performance

The PFLISA architecture was designed to maximize the cytokine coverage across a wide concentration range while minimizing sample consumption. Capture antibodies specific to TNF-α, IL-1α, IL-1β, IL-2, IL-4, IL-5, IL-6, IL-8, IL-10, IL-12p70, IL-13, and IFN-γ were spatially encoded in a 7 × 7 array at the bottom of each well of a 96-well plate (Fig. 1(a)). Each cytokine was represented by three spatially distributed spots, creating three independent measurements per analyte while retaining all 12 targets within a single well. The spacing between replicate spots reduced sensitivity to localized artifacts such as dust, scratches, or debris (Fig. S1). The first two rows of the plate were reserved for standard curves, with the starting concentration for each cytokine provided in Table S1. For each test sample, 15 μL of plasma from a healthy donor or patient was diluted four-fold to a final volume of 60 μL and loaded into one well according to the plate map in Fig. S2.

**Fig. 1.**
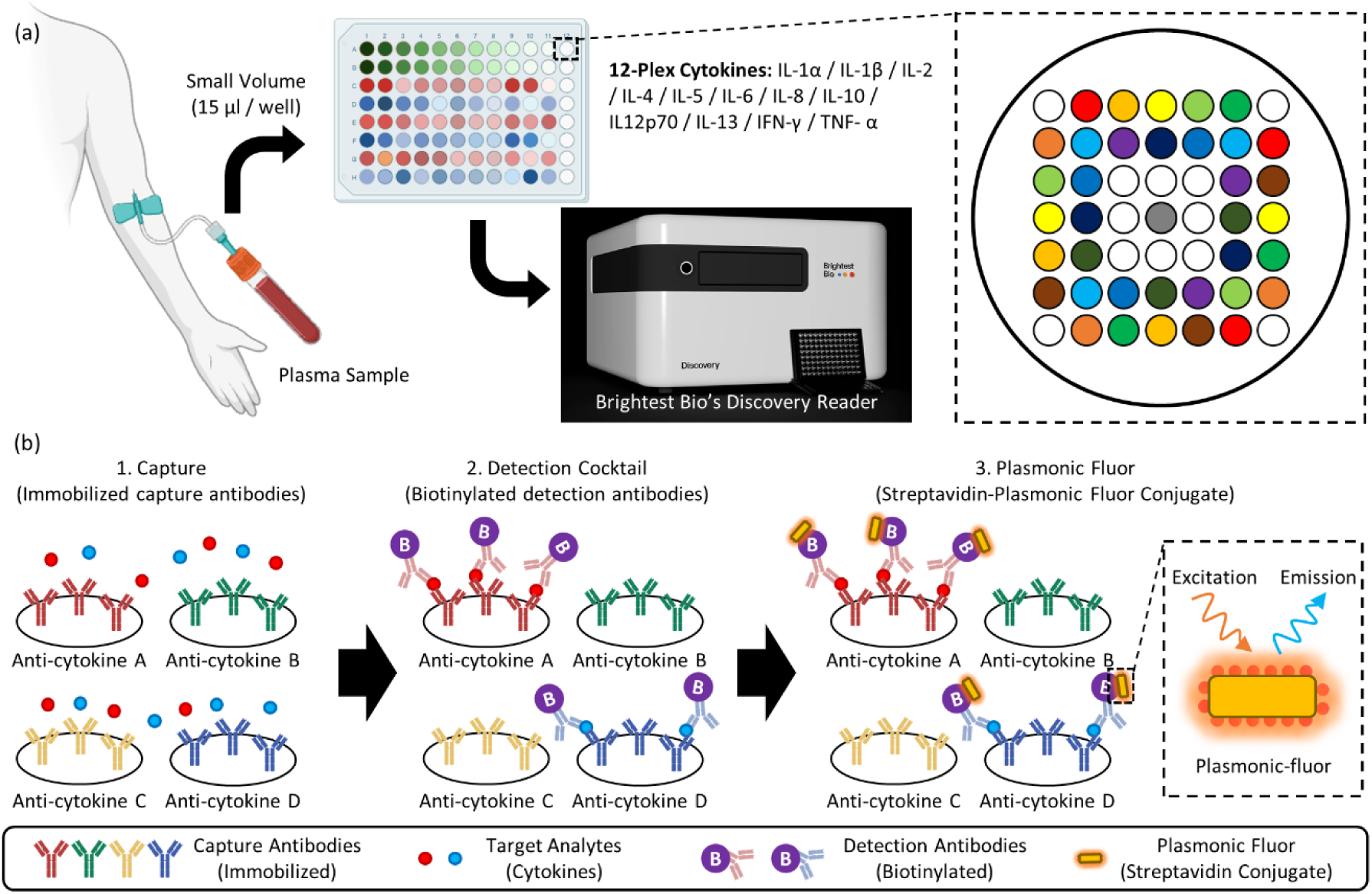
Multiplexed PFLISA platform and assay workflow. (a) Schematic illustration of the multiplexed PFLISA platform. (b) Stepwise workflow for cytokine quantification within a single well of a 96-well plate.

Spatial multiplexing was paired with Plasmonic Fluor signal amplification in a sandwich immunoassay format (Fig. 1(b)). After cytokines bound to their corresponding capture antibodies, detection antibodies and Plasmonic Fluor labels generated a fluorescence signal for each arrayed target. Fluorescence intensities were quantified using the Brightest Bio analysis software. As shown in Fig. S3, serial dilutions of the standards produced monotonically decreasing fluorescence with increasing dilution, confirming the expected concentration-dependent response across the multiplexed array. This combination of spatial multiplexing and signal amplification was designed to extend quantification into the sub-pg/mL range while measuring all 12 panel cytokines in parallel from the same sample.

To minimize the influence of experimental artifacts, median values from the triplicates were used for subsequent analysis. Standard curves for each of the 12 cytokines were generated from serial dilutions of known concentrations (Fig. 2(a–l)). Curve fitting was performed using either four-parameter or five-parameter logistic regression, with the optimal model selected based on goodness-of-fit for each cytokine. The lower limit of detection (LOD) was defined as the blank signal plus three times the standard deviation of the blank (3σ). Specifically, the quantifiable range was defined by calibration points meeting predefined accuracy and precision criteria. The LLOQ and ULOQ were assigned the lowest and highest concentrations, respectively, at which both accuracy (% bias) and precision (%CV) were within 25%; all intermediate points were required to remain within 20% for both bias and CV. A calibration curve was considered valid when more than 75% of points within the dynamic range had a total error (%bias + %CV) below 40%. Each cytokine exhibited an analyte-specific quantitative range spanning multiple orders of magnitude. Maximum detectable concentrations extended up to the highest tested standards for TNF-α (6000 pg/mL), IL-1β (2000 pg/mL), IL-4 (200 pg/mL), IL-8 (500 pg/mL), IL-10 (400 pg/mL), IL-12p70 (800 pg/mL), and IL-13 (600 pg/mL). In contrast, ULOQs were reached for IFN-γ (1174.31 pg/mL), IL-1α (249.16 pg/mL), IL-2 (607.48 pg/mL), IL-5 (185.37 pg/mL), and IL-6 (138.19 pg/mL). The LLOQs achieved ranged from 3.4 to 110 fg/mL across the 12 cytokines, highlighting the high sensitivity of the assay. Detailed LLOQ values for individual cytokines are summarized in Table S2.

**Fig. 2.**
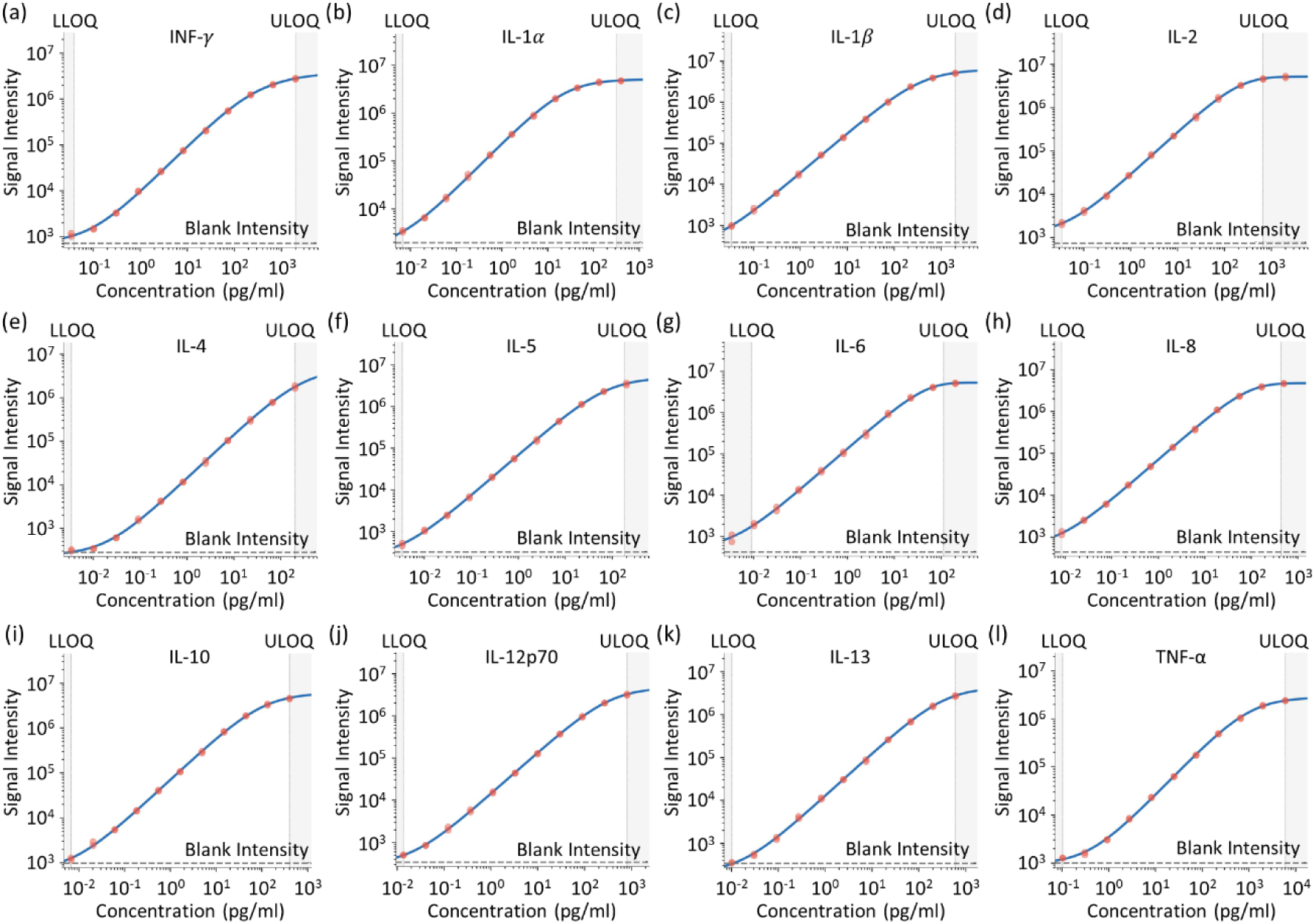
Calibration curves for the 12 target cytokines. (a–l) Standard calibration curves for the 12 target cytokines.

Compared with reported conventional multiplex cytokine assays, including Luminex (*30, 31, 32, 33*), cytometric bead array (CBA) (*34, 35*), and mesoscale discovery (MSD) platforms (*32, 33, 34, 35, 36*), PFLISA achieved substantially lower LLOQs for the cytokines evaluated in this study (Table S2). For several targets, the LLOQs were comparable to or lower than values reported for the single-molecule array (Simoa), one of the most sensitive available methods (*11, 37, 38*). Importantly, this level of sensitivity was achieved within a 12-plex, single-well format, allowing cytokines spanning markedly different concentration ranges to be measured together. The resulting analytical coverage extended into basal concentrations observed in healthy individuals while retaining quantification of highly elevated inflammatory mediators in sepsis. Thus, the ultrasensitivity expanded the portion of the cytokine network that remains quantitatively observable, bringing low-abundance regulatory signals into the same patient-level profile as abundant inflammatory mediators without partitioning limited samples across separate assays.

The PFLISA 12-plex calibrators were also compared with corresponding NIBSC/WHO cytokine reference standards (Table S3). Across the 12 cytokines, the ratio of the NIBSC/WHO standard concentration to the PFLISA standard concentration ranged from 0.8 to 1.2, with a median ratio of 1.0; 10 of 12 analytes had ratios between 0.9 and 1.1. This close alignment provides an external reference for the assay calibration and supports confidence in the quantitative concentrations reported across the multiplex panel.

Within each well, three replicate spots per cytokine provided technical replicates and an internal measure of assay precision. The coefficient of variation (CV) across replicate spots averaged 5.7% (Fig. S4), demonstrating low within-well variability across the multiplexed array. Although certain cytokines at specific concentrations exhibited CVs greater than 10%, all values remained below the 15% threshold recommended by the Clinical and Laboratory Standards Institute (CLSI EP05-A3) for clinically acceptable precision (*39*).

Precision was further evaluated across 16 independent plates at low, medium, and high QC concentrations (Table S4). At medium and high QC levels, average intraplate CVs ranged from 3.4% to 8.6%, while interplate CVs ranged from 8.9% to 15.0% across all 12 cytokines. At the low QC level, 11 of 12 cytokines exhibited intraplate CVs of 5.5–14.6% and interplate CVs of 9.1–18.9%. The exception was IL-1α at 0.02 pg/mL, which showed greater variability (35.4% intraplate and 36.8% interplate). Thus, 35 of 36 analyte-by-QC conditions demonstrated both intra- and interplate CVs below 20%, with variability increasing for IL-1α at the lowest tested concentration.

Matrix spike-recovery studies further characterized assay performance in plasma across low, medium, and high cytokine concentrations (Table S5). Mean plasma recovery ranged from 58.5% to 112.1% across the 36 analyte-by-concentration conditions, with 35 of 36 mean recoveries falling between 60% and 120%. Recovery was analyte dependent, with the lowest mean recovery observed for low-level IL-1α. Corresponding serum spike-recovery measurements are provided in Table S6.

### Patient Characteristics

A total of 100 patients with sepsis and 97 healthy controls were included in this study. The demographic and clinical characteristics of the study populations are summarized in Table 1, with detailed individual clinical and physiological characteristics of the sepsis cohort provided in Table S7. The mean age was 60.3 ± 16.0 years, with 57 males (57.0%). Gram-positive bacteria accounted for the majority of infections (58.0%), followed by Gram-negative bacteria (39.0%). The four most commonly isolated pathogens were *Escherichia coli* (20.0%), coagulase-negative staphylococci (CoNS, 20.0%), *Staphylococcus aureus* (14.0%), and *Klebsiella pneumoniae* (9.0%). The overall in-hospital mortality rate was 33.0% (33/100).

**Table 1.** Patient demographics for healthy and septic individuals in the study. The “Other” race category included patients recorded as mixed race, or not reported in the source clinical data. The “Other” pathogen category included cases with mixed/polymicrobial, or fungal etiologies.

|  | **Healthy (n=97)** | **Patient (n = 100)** |
|---|---|---|
| Age (years) | 47.3 (21-81) | 60 (20-92) |
| Sex, Male | 46 (47.4%) | 57 (57.0%) |
| Race |  |  |
| Asian | 4 (4.1%) | 27 (27.0%) |
| Black | 23 (23.7%) | 3 (3.0%) |
| White | 70 (72.2%) | 36 (36.0%) |
| Others | - | 34 (34.0%) |
| Type of Pathogen |  |  |
| Gram Positive | - | 58 (58.0%) |
| Gram Negative | - | 39 (39.0%) |
| Others | - | 3 (3.0%) |
| Mortality | - | 33 (33.0%) |

When stratified by survival status, non-survivors demonstrated significantly lower platelet counts than survivors (p = 0.003) and were more likely to have received prior antibiotic treatment (p = 0.038). Organ dysfunction severity also differed significantly between groups (p = 0.004), with multi-organ dysfunction present in 41.9% of non-survivors compared with 16.7% of survivors. No significant differences in age, sex, vital signs, or pathogen type were observed between survivors and non-survivors.

### Cytokine Profiles in Sepsis Versus Healthy Controls

The assay’s combined sensitivity and multiplexing enabled a multi-cytokine comparison in patients with sepsis and healthy controls (**Table S8**), including mediators that differed greatly in absolute concentration. Of the 12 cytokines analyzed, 11 were significantly elevated in septic patients after FDR correction (all FDR-adjusted p < 0.001). IL-12p70 was the only cytokine that did not differ significantly between groups (FDR-adjusted p = 0.801).

Among pro-inflammatory cytokines, IL-6 and IL-8 exhibited the largest elevations, with median concentrations approximately 78.5-fold and 52.3-fold higher in septic patients than in healthy controls, respectively (Fig. 3). IFN-γ, IL-1β, and IL-2 also demonstrated marked increases (8.6– 11.7-fold). Among anti-inflammatory cytokines, IL-10 demonstrated the greatest elevation (10.8-fold), whereas IL-4, IL-5, and IL-13 were elevated to a lesser extent (2.1–2.6-fold).

**Fig. 3.**
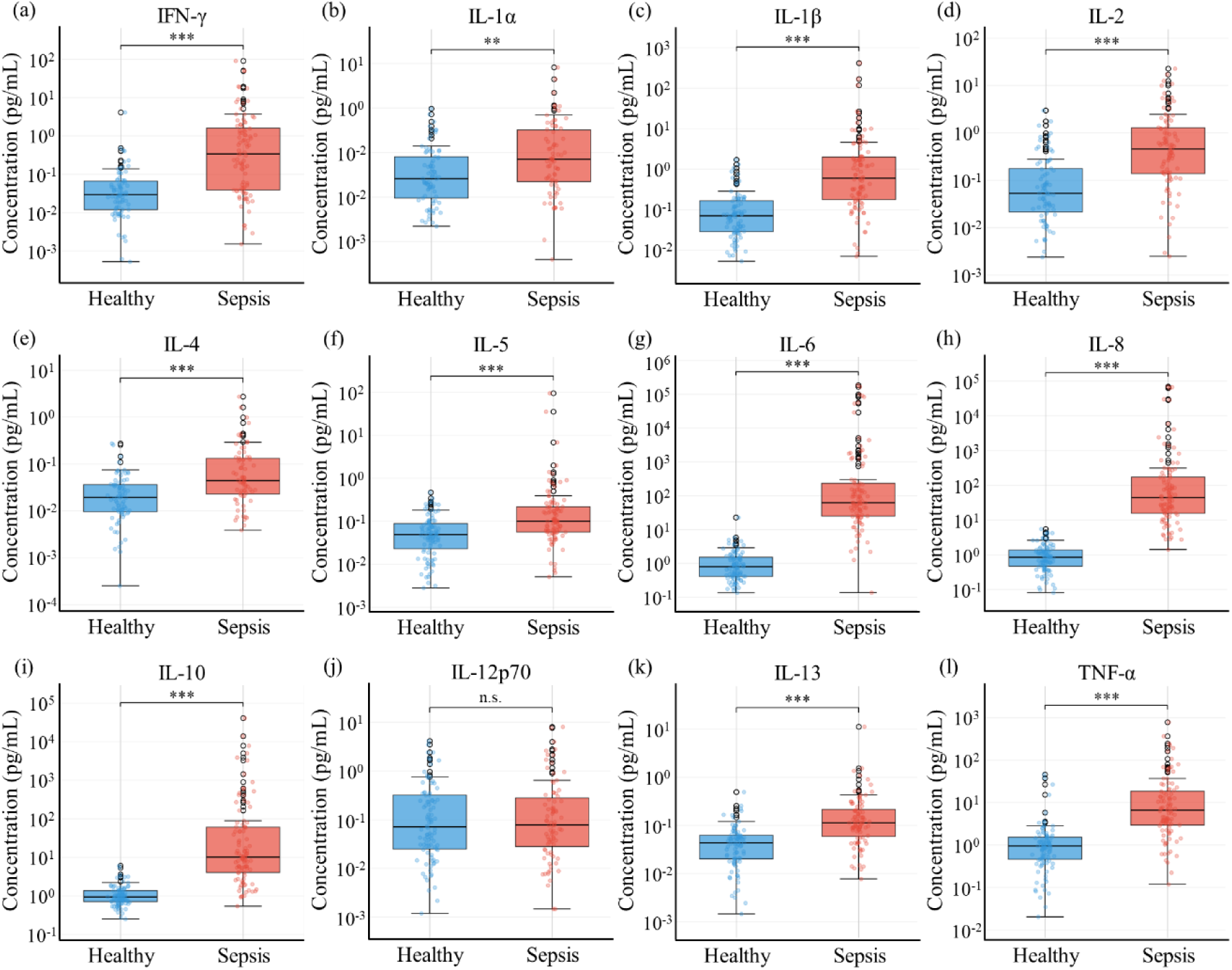
Cytokine concentrations in healthy controls and sepsis patients. Comparison of measured cytokine concentrations between healthy control (blue, n = 97) and septic patients (red, n = 100) for all 12 target cytokines measured by PFLISA (a-l). Boxes represent the interquartile range (IQR; 25th-75th percentiles), with the solid black horizontal line within each box indicating the median. Whiskers extend to 1.5×IQR from the box edges, and data points beyond the whiskers are plotted as outliers (open circles). Individual measurements are overlaid as semi-transparent dots. Statistical significance between the two groups is indicated by asterisk; **p < 0.01, *** p<0.001, and n.s. not significant.

To assess the overall pro-inflammatory versus anti-inflammatory cytokine balance, composite scores were calculated using the geometric mean of available cytokine concentrations within each panel. The pro-inflammatory composite score was 10.6-fold higher in sepsis patients than in healthy controls (p < 0.001), while the anti-inflammatory composite score was elevated 3.9-fold (p < 0.001). The Pro/Anti ratio was 2.0-fold higher in sepsis patients (p < 0.001), indicating concurrent elevation of both cytokine groups, with a greater relative increase in the pro-inflammatory component.

Because all 12 mediators were measured from the same low-volume sample, the data could also be examined as integrated patient-level cytokine profiles rather than separate single-analyte comparisons. The heatmaps in Fig. S5 show a marked contrast between groups: patients with sepsis displayed heterogeneous, broadly elevated profiles, particularly for IL-6, IL-8, TNF-α, and IL-10, whereas healthy controls exhibited uniformly low concentrations across the panel. Simultaneous measurement across this concentration range preserved both dominant inflammatory signals and lower-abundance components within each patient’s profile, revealing substantial inter-individual heterogeneity beyond that apparent from group-level comparisons alone.

### Cytokine Profiles by Pathogen Type

Cytokine concentrations were compared between patients with Gram-positive (n = 58) and Gram-negative (n = 39) bacterial infections, excluding mixed (n = 2) and fungal (n = 1) infections (Table S9). Four cytokines were significantly elevated in Gram-negative compared with Gram-positive infections after FDR correction: IL-6 (FDR-adjusted p = 0.022), IL-8 (FDR-adjusted p = 0.022), TNF-α (FDR-adjusted p = 0.022), and IL-10 (FDR-adjusted p = 0.022) (Fig. 4). Of these, three were pro-inflammatory (IL-6, IL-8, TNF-α) and one was anti-inflammatory (IL-10). Thus, the Gram-negative profile was characterized by coordinated elevation of three pro-inflammatory mediators together with the regulatory cytokine IL-10.

**Fig. 4.**
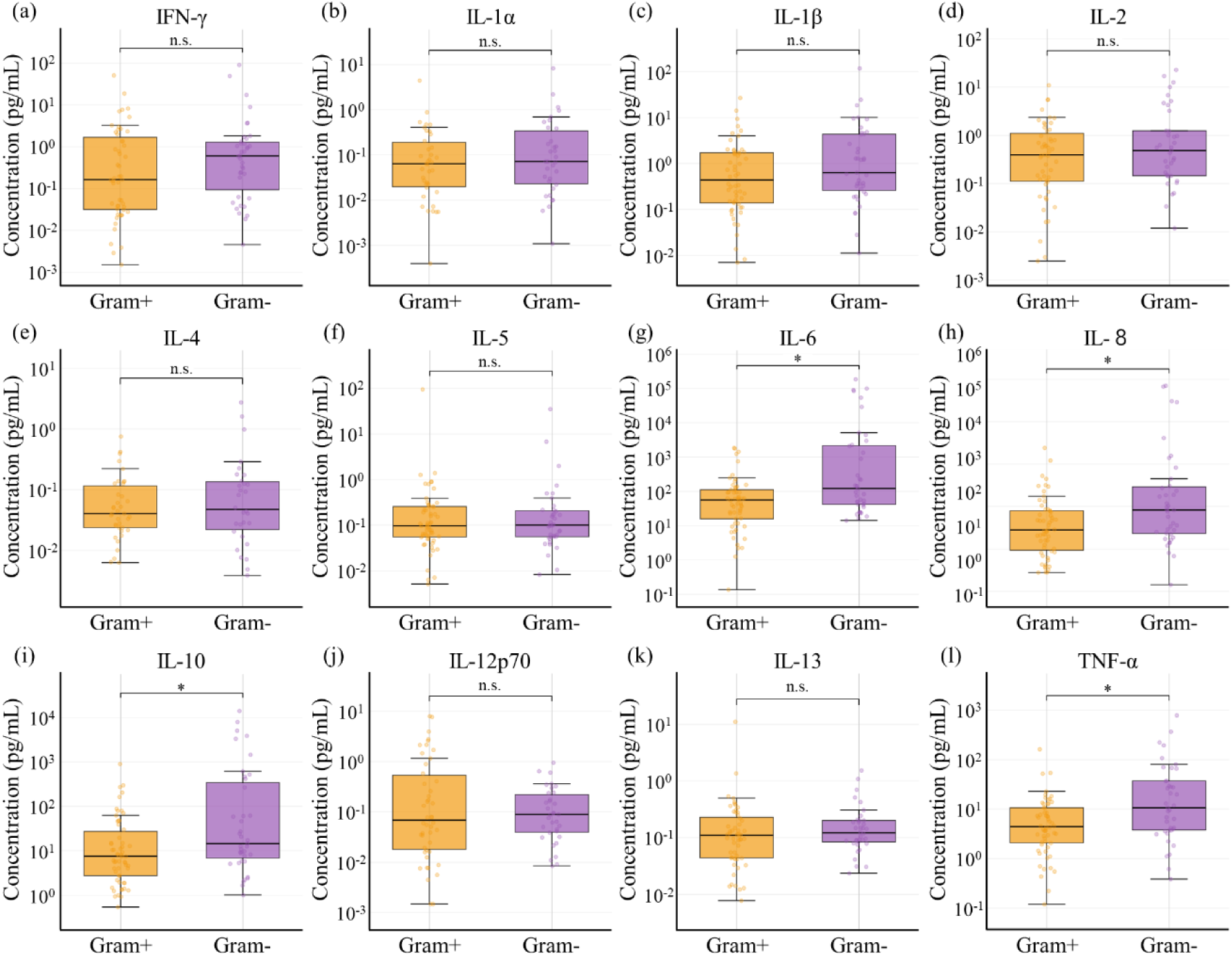
Cytokine concentrations by bacterial Gram classification. Comparison of measured cytokine concentrations with gram positive (orange, n = 58) and gram negative (purple, n = 39) bacterial infections among sepsis patients for all 12 target cytokines. Boxes represent the interquartile range (IQR; 25th-75th percentiles), with the horizontal line within each box indicating the median. Whiskers extend to 1.5×IQR from the box edges. Individual measurements are overlaid as semi-transparent dots. Statistical significance was assessed by Mann-Whitney U test with Benjamini-Hochberg FDR correction for 12 simultaneous comparisons. *p < 0.05; n.s., not significant.

The pro-inflammatory composite score was higher in Gram-negative than in Gram-positive infections (p = 0.030). The core Pro/Anti ratio, computed from IL-6, IL-8, and TNF-α (pro-inflammatory) and IL-5 and IL-10 (anti-inflammatory), was significantly higher in Gram-negative infections (p = 0.002), indicating greater pro-relative increase in the pro-inflammatory component despite concurrent elevation of IL-10.

A comparison across the four most common pathogens (*E. coli*, *S. aureus*, CoNS, and *K. pneumoniae*) revealed trends consistent with the Gram-positive versus Gram-negative findings. CoNS-associated infections exhibited consistently lower cytokine concentrations across most analytes. Although no individual cytokine reached FDR-adjusted significance in the four-group comparison, exploratory post-hoc analyses revealed significant pairwise differences between CoNS and *E. coli* for IFN-γ and IL-6.

### Cytokine profiles by survival status

Cytokine concentrations stratified by survival status are presented in Table S10. No individual cytokine reached statistical significance after FDR correction. Nevertheless, IL-6 and IL-8, both pro-inflammatory cytokines, demonstrated nominally significant elevations in non-survivors (raw p = 0.011 for both). Conversely, IFN-γ was nominally higher in survivors (raw p = 0.030), suggesting a potential divergence in immune response patterns associated with survival.

The core Pro/Anti ratio demonstrated a trend toward higher values in non-survivors compared with survivors (p = 0.063), consistent with greater pro-inflammatory predominance in patients who did not survive.

### Cytokine profiles by organ dysfunction severity

Cytokine concentrations across organ dysfunction severity groups are presented in **Table S11**. Among 85 patients with complete data for organ dysfunction scoring, 32 (37.6%) had no dysfunction, 31 (36.5%) had single-organ dysfunction, and 22 (25.9%) had multi-organ dysfunction.

No individual cytokine reached FDR-adjusted significance. Nominally significant trends were observed for IL-2 (raw p = 0.011), IL-8 (raw p = 0.010), and IL-10 (raw p = 0.036), with concentrations increasing with the number of dysfunctional organs. Post-hoc analysis for IL-8 identified significant pairwise differences between the no-dysfunction group and both the single-organ (p = 0.025) and multi-organ (p = 0.018) groups, showing progressively higher concentrations across dysfunction categories.

### Cytokine Correlations

Simultaneous measurement of the 12 cytokines also enabled analysis of coordination among network components. Spearman correlation analysis in sepsis patients identified 13 cytokine pairs with strong correlations (|ρ| > 0.5, p < 0.05) (Fig. 5). The strongest correlation was between IL-6 and IL-10 (ρ = 0.821, p < 0.001), linking the most markedly elevated pro-inflammatory and anti-inflammatory cytokines. Other strong relationships included IL-1β with IL-6 (ρ = 0.701), TNF-α (ρ = 0.656), IL-8 (ρ = 0.567), and IL-10 (ρ = 0.689); IL-6 with IL-8 (ρ = 0.685) and TNF-α (ρ = 0.667); and TNF-α with IL-10 (ρ = 0.736). These patterns show that pro-inflammatory activation was not isolated but was coordinated with both other inflammatory mediators and the anti-inflammatory response represented by IL-10.

**Fig. 5.**
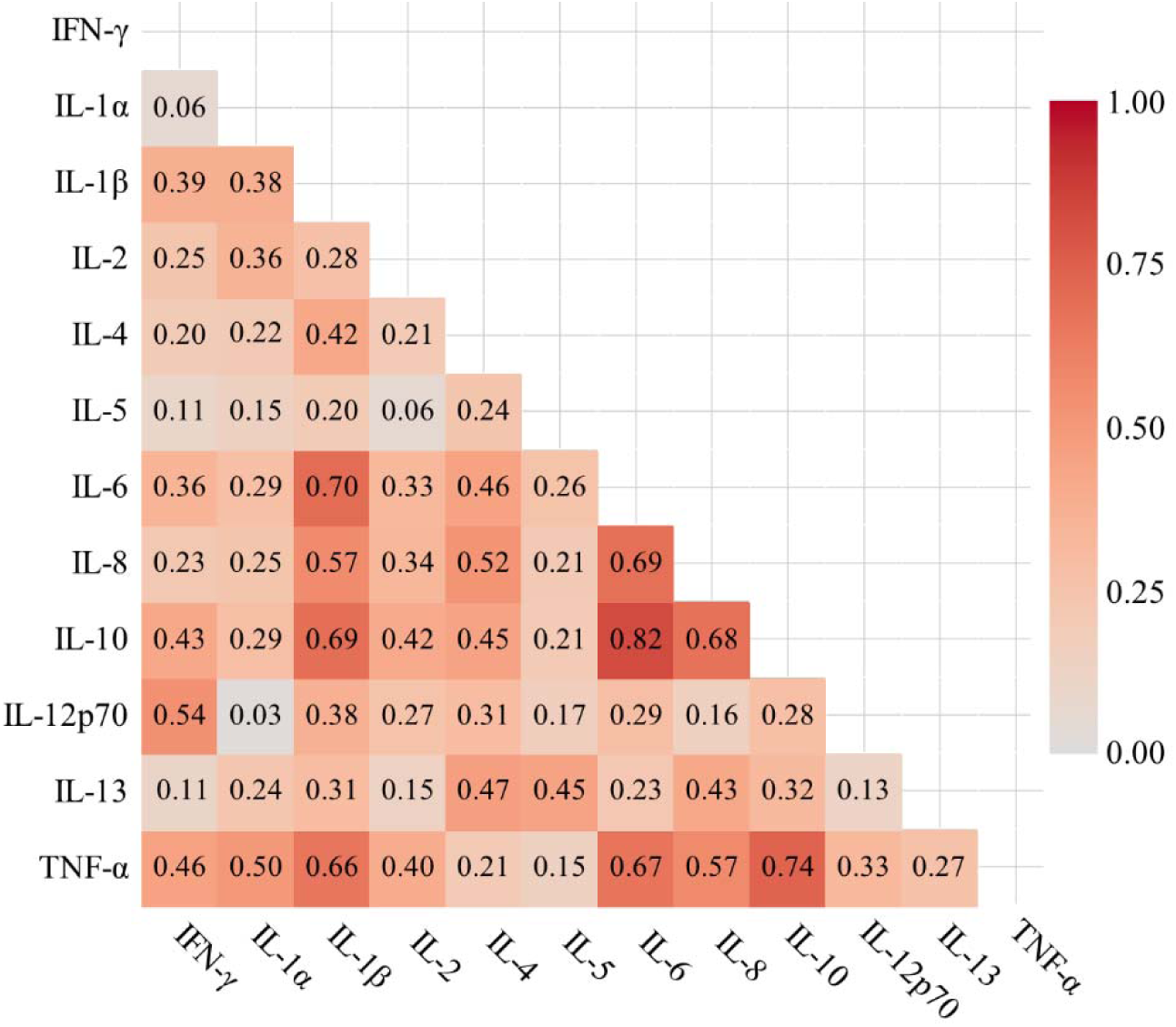
Cytokine correlation matrix in sepsis patients.

## DISCUSSION

The principal technological advance demonstrated here is not sensitivity alone, but the ability to extend quantitative measurement across cytokines spanning markedly different concentration ranges in the same sample. By combining femtomolar detection, 12-plex measurement, broad concentration coverage, low sample input, and a standard plate-based format, PFLISA expands the portion of the immune response that remains quantitatively observable rather than restricting analysis to its most abundant mediators. Applying the platform to 100 sepsis patients and 97 healthy controls, we detected significant elevations in 11 of 12 cytokines, resolved disproportionate pro-inflammatory activation (Pro/Anti ratio 2.0-fold higher than controls, p < 0.001), identified pathogen-associated differences, and mapped coordinated relationships among cytokines. These findings demonstrate that expanded analytical coverage allows for low-abundance regulatory signals to be interpreted in the same patient-level profile as highly elevated inflammatory markers. The sepsis cohort therefore serves as a biological demonstration of the platform’s capacity to measure an immune system as a network rather than as a collection of isolated analytes.

A central biological finding was the concurrent elevation of both pro-inflammatory and anti-inflammatory cytokines, with a substantially greater increase in the pro-inflammatory component. The pro-inflammatory composite score was 10.6-fold higher than in healthy controls, compared with a 3.9-fold increase in the anti-inflammatory composite, yielding a Pro/Anti ratio approximately twice that of healthy individuals. Although sepsis has historically been described as an initial cytokine storm followed by compensatory anti-inflammatory response syndrome (CARS) (*4, 25*), our cross-sectional data support a model in which these responses are concurrently active and quantitatively coupled, with the pro-inflammatory component predominating. Our cross-sectional measurements are consistent with this concurrent activation. Both cytokine groups were elevated, but the magnitude of increase was greater for the pro-inflammatory components. Resolving this relative balance depended on quantitative measurement of both highly elevated inflammatory cytokines and substantially lower-abundance regulatory mediators within the same sample.

The strong positive correlation between IL-6 and IL-10 (ρ = 0.821) provides additional support for this coordinated inflammatory and regulatory responses. IL-6, a principal mediator of the acute inflammatory response in sepsis, is known to stimulate IL-10 release through STAT3-dependent signaling (*40, 41*). The tight coupling between these two cytokines observed in our data suggests that the compensatory anti-inflammatory response is directly proportional to the magnitude of the pro-inflammatory stimulus. Nevertheless, the Pro/Anti ratio exceeding 2.0 in sepsis patients indicates that this compensatory mechanism is insufficient to fully counterbalance the pro-inflammatory burden, thereby resulting in a net pro-inflammatory state.

Prior ELISA- and Luminex-based studies have reported elevated IL-6, IL-8, and TNF-α in sepsis (*42, 43*), consistent with our findings. However, regulatory cytokines such as IL-4, IL-5, and IL-13 occur at much lower concentrations and are below the quantification limits of conventional multiplex assays (*44*). The value of Plasmonic Fluor enhancement in this setting is therefore not only the detection of lower concentrations, but expansion of the portion of the cytokine network that remains quantitatively observable. By bringing low-abundance regulatory mediators into the same measurable profile as abundant inflammatory cytokines, the platform preserves information across a wider concentration spectrum. This expanded coverage enabled the composite balance and correlation analyses that form the central biological interpretation of this study.

Gram-negative infections were associated with significantly higher concentrations of IL-6, IL-8, TNF-α, and IL-10 (all FDR-adjusted p = 0.022) compared with Gram-positive infections. These differences likely reflect the distinct molecular mechanisms by which Gram-negative and Gram-positive bacteria activate innate immunity. Lipopolysaccharide (LPS), a component of the Gram-negative bacterial outer membrane, is a potent activator of Toll-like receptor 4 (TLR4) signaling, which triggers a robust pro-inflammatory cascade including NF-κB activation and downstream cytokine production (*45*). In contrast, Gram-positive bacteria primarily activate immune responses through lipoteichoic acid and peptidoglycan via TLR2, which generally elicits a less pronounced cytokine response (*46*). The core Pro/Anti ratio was significantly higher in Gram-negative infections (p = 0.002), indicating that the greater pro-inflammatory burden in Gram-negative sepsis was not fully compensated by the accompanying anti-inflammatory response. Among the four most isolated pathogens, CoNS-associated infections consistently exhibited the lowest cytokine concentrations across most analytes. This observation is consistent with the nature of CoNS as an opportunistic pathogen with lower virulence, which may provoke a less intense inflammatory response compared with more pathogenic organisms such as *E. coli* or *K. pneumoniae* (*22, 47*).

Although no individual cytokine comparison reached FDR-adjusted significance when stratified by survival status, several findings merit consideration. IL-6 and IL-8 were nominally elevated in non-survivors (raw p = 0.011), consistent with previous reports identifying IL-6 as an independent predictor of sepsis mortality (*43*). These cytokines are key mediators of the acute-phase response and neutrophil chemotaxis, and their sustained elevation may reflect ongoing or uncontrolled infection. An unexpected observation was the nominally higher IFN-γ concentration in survivors compared with non-survivors. IFN-γ is a Th1 cytokine critical for macrophage activation and intracellular pathogen clearance (*48*). Higher IFN-γ levels in survivors may indicate preserved adaptive immune function, which could contribute to more effective pathogen elimination. This finding warrants further investigation in prospective cohorts. The core Pro/Anti ratio demonstrated a trend toward higher values in non-survivors (p = 0.063), suggesting that the degree of pro-inflammatory/anti-inflammatory imbalance may be relevant to clinical outcomes. Similarly, IL-2, IL-8, and IL-10 exhibited nominally significant increases with organ dysfunction severity. The absence of statistical significance after FDR correction may reflect limited statistical power inherent in the sample size of this study. These findings should be considered hypothesis-generating and require validation in larger, prospective investigations.

Viewed as an analytical system, PFLISA provides high information density per unit of sample. Its LLOQs of 3.4–110 fg/mL represented a 100–1000-fold sensitivity improvement over standard Luminex and CBA platforms, while the spatially encoded array measured 12 cytokines in triplicate from the same well. Precision across 16 plates was strongest at medium and high QC concentrations, where interplate CVs remained ≤15% across all analytes, while performance at the lowest concentrations was more analyte dependent. Plasma spike-recovery measurements further demonstrated quantitative recovery across a broad concentration range, with mean recoveries of 58.5–112.1% across analytes and spike levels. This combination of these capabilities is critical: sensitivity made sub-pg/mL cytokines such as IL-4, IL-5, and IL-13 quantifiable, multiplexing placed them in context with abundant IL-6, IL-8, and TNF-α, and the 15 μL input preserved limited clinical material. The platform thus converts analytical sensitivity into broader biological visibility.

Operationally, the 12-analyte profile is generated in parallel from 15 μL of plasma rather than through serial single-analyte tests. The 96-well format and established sandwich immunoassay chemistry support batch processing and integration with familiar laboratory workflows. These features increase information yield per sample and per assay run, an important consideration for immune monitoring when specimen volume, time, and laboratory capacity are constrained. Further automation and prospective validation could extend this approach from research profiling toward longitudinal and clinically actionable immune-state measurement. Additionally, the underlying Plasmonic Fluorescence detection strategy may also provide a path towards incorporation into more automated laboratory diagnostics workflows.

This study has several limitations. First, the cross-sectional design captured cytokine concentrations at one time point and therefore could not resolve how network states change during sepsis progression. Serial sampling will be important for testing whether the platform can identify transitions in inflammatory balance within individual patients. Second, although the cohort supported the primary sepsis-control comparison, statistical power was limited for survival and organ-dysfunction subgroups. Third, the organ dysfunction score was derived from four available clinical parameters and was not a complete SOFA assessment because Glasgow Coma Scale, creatinine, and FiO2 data were unavailable. Fourth, the 12-cytokine panel captures a highly informative but necessarily incomplete portion of the broader immune network. Additionally, mediators and longitudinal measurements will be needed to more fully characterize immune-state dynamics. Finally, this study establishes analytical and biological proof of concept but does not define clinical decision thresholds or a fully automated turnaround time. Multicenter studies with serial sampling, larger cohorts, complete clinical scoring, and workflow validation will be needed to establish clinical utility.

This study establishes PFLISA as an information-rich immune-profiling platform that combines femtomolar sensitivity, 12-plex measurement, broad dynamic range, and 15 µL sample input within a single-well, 96-well format. This analytical advancement brings low-abundance regulatory cytokines into the same profile as abundant inflammatory mediators, enabling assessment of inflammatory balance, patient heterogeneity, pathogen-associated responses, and cytokine coordination without requiring new assays for each question. Applied to sepsis, the platform revealed concurrent but disproportionate pro-inflammatory activation and demonstrated how ultrasensitive multiplexing can convert limited clinical samples into system-level immune information.

## MATERIALS AND METHODS

### Plasmonic-Enhanced Fluorescence Assay

A commercial Human Inflammation 12-plex Kit (HIF-12A, Brightest Bio, Missouri, USA) was used for multiplexed cytokine detection. The kit comprises a 96-well plate with antibodies against 12 cytokines including TNF-α, IL-1α, IL-1β, IL-2, IL-4, IL-5, IL-6, IL-8, IL-10, IL-12p70, IL-13, and IFN-γ, immobilized in a 7×7 array format within each well.

Each cytokine is represented by three spatially distributed spots within the 7×7 array, as illustrated in Fig. 1 and Fig. S1. A positive control was positioned at the center of the array to confirm assay performance and facilitate array alignment. Blank spots were placed around the central region and at the four corners to minimize potential interference from the positive signals and to minimize edge-associated artifacts.

### Human Inflammation 12-plex (HIF-12A) PFLISA

To establish standard curves for the 12 cytokines, lyophilized cytokine cocktail solutions were reconstituted by adding 500 μL of sample diluent (Brightest Bio, St. Louis, MO) to get the standard 1 concentration as listed in Table S1. 150 μL of standard 1 was then mixed with 300 μL of sample diluent for a three-fold serial dilution. This process was repeated 10 times, to generate a total of 11 standard concentrations.

Corresponding NIBSC/WHO standard codes and the ratio of NIBSC/WHO standard concentration to PFLISA 12-plex standard concentration are provided in Table S3.

Patient plasma samples were aliquoted into 50 μL portions and centrifuged at 10,000 x g for 10 minutes to remove cellular debris. A 4-fold diluted plasma sample was prepared by adding 45 μL of the supernatant to 135 μL of sample diluent. The prepared standard samples and diluted plasma samples were dispensed into the 12-plex 96-well plate, with each well receiving 60 μL, as indicated in Fig. S2. Each sample well contained the equivalent of 15 µL of undiluted plasma. The plate was then incubated at 650 rpm for 3 hours on an orbital microplate shaker (Southwest Science, SBT 1500, Roebling, NJ) to allow cytokines to bind to the immobilized antibodies.

After incubation, unbound proteins were removed by washing the plate three times with Brightest Bio 1x wash buffer. Subsequently, 12-plex detection antibody cocktail was reconstituted in detection antibody diluent. 50 μL of detection antibody solution per well, was added and the plates were incubated at 650 rpm for one hour. The plate was washed three times with wash buffer before adding reconstituted PF solution at 50 μL per well. The plate was incubated in the dark for 20 minutes. Following six additional washes with wash buffer, 200 μL of wash buffer was added to each well, and the fluorescence intensity was measured using Brightest Bio’s Discovery reader.

### Analytical Performance Characterization

Analytical precision was evaluated across 16 plates using low-, medium-, and high-concentration quality-control (QC) samples for each of the 12 cytokines. Average intraplate and interplate coefficients of variation (%CV) were calculated at each QC concentration. Matrix spike recovery was additionally evaluated at low, medium, and high spike concentrations in serum (N = 5) and plasma (N = 4). Percent recovery was calculated relative to the known amount of cytokine added to each matrix.

### Software and Statistical Analysis

Fluorescence images were acquired using the Brightest Bio Discovery reader, and fluorescence intensities were quantified using the accompanying Empower analysis software. Using the Brightest Bio data analysis suite Empower at https://app.brightestbio.com/, the template for the given plate map (Fig. S2) was created and the data was analyzed using that template. Four-parameter or five-parameter logistic regression models (4PL or 5PL) fitted standard curves were generated by the software and back-calculated concentrations for the samples was provided.

Cytokine concentrations were reported as medians with interquartile ranges (IQR), as distributions were right-skewed and non-normal (Shapiro–Wilk test). Age was reported as mean ± standard deviation (SD) or median [IQR], depending on normality. Categorical demographic and clinical variables, including sex, race, type of pathogen, and mortality were reported as counts with percentages (Table 1).

For comparisons of cytokine concentrations between two groups (septic patients versus healthy controls; Gram-positive versus Gram-negative infections; survivors versus non-survivors), the Mann–Whitney U test was employed. For comparisons across three or more groups (organ dysfunction severity; major infection sources), the Kruskal–Wallis test was applied, followed by Dunn’s post-hoc test with Bonferroni adjustment when overall significance was detected. To account for multiple testing across 12 cytokines within each comparison, p-values were adjusted using the Benjamini–Hochberg false discovery rate (FDR) correction. Two-sided FDR-adjusted p < 0.05 was considered statistically significant for individual cytokine comparisons.

To assess the balance between pro-inflammatory and anti-inflammatory cytokine responses, composite scores were calculated. The 12 measured cytokines were classified as pro-inflammatory (IFN-γ, IL-1α, IL-1β, IL-2, IL-6, IL-8, IL-12p70, and TNF-α) or anti-inflammatory (IL-4, IL-5, IL-10, and IL-13) based on established functional roles (*11, 49*). This grouping was used as a summary framework and does not imply that individual cytokines have exclusively pro- or anti-inflammatory activity. For each patient, pro-inflammatory and anti-inflammatory composite scores were computed as the geometric mean of available cytokine concentrations within each panel (minimum two cytokines required). The pro-inflammatory/anti-inflammatory (Pro/Anti) ratio was defined as the pro-inflammatory composite score divided by the anti-inflammatory composite score. A core Pro/Anti ratio was additionally computed using the three pro-inflammatory cytokines (IL-6, IL-8, and TNF-α) and two anti-inflammatory cytokines (IL-5 and IL-10) with the lowest missing data rates. Composite scores were compared between groups using the Mann–Whitney U test or Kruskal–Wallis test as appropriate. FDR correction was not applied to composite score comparisons, as these represented pre-specified summary measures rather than multiple individual tests.

An organ dysfunction score (range 0–4) was derived from four SOFA-based criteria: hepatic dysfunction (bilirubin ≥ 1.2 mg/dL), thrombocytopenia (platelets < 150 × 10³/μL), cardiovascular dysfunction (mean arterial pressure < 70 mmHg), and respiratory dysfunction (respiratory rate ≥ 22/min). Patients were categorized as having no dysfunction (score = 0), single-organ dysfunction (score = 1), or multi-organ dysfunction (score ≥ 2). Patients with missing values for any component variable were excluded from this analysis.

Spearman rank correlation coefficients were calculated to assess pairwise relationships among the 12 cytokines in sepsis patients. Correlations with |ρ| > 0.5 and p < 0.05 were considered strong.

Additional exploratory subgroup analyses were performed by prior antibiotic treatment, clinical severity category (derived from medical record text), infection site, comorbidity status, modified systemic inflammatory response syndrome (SIRS) criteria, thrombocytopenia, and hepatic dysfunction. These analyses were designated as exploratory and interpreted accordingly.

Effect sizes for two-group comparisons were reported as fold change (ratio of group medians) and rank-biserial correlation coefficient (r). All analyses were performed using Python 3.x with SciPy, statsmodels, and scikit-posthocs packages.

### Samples

Plasma samples from 97 healthy individuals were purchased from Discovery Life Sciences (California, USA). These normal plasma samples were collected from healthy donors without known disease. Blood was drawn into EDTA-treated tubes and centrifuged at 2,000 g for 15 minutes at 4°C to obtain platelet-free plasma.

Patient plasma samples were obtained from 100 individuals presenting with sepsis at the University of California, Irvine Medical Center, a tertiary academic medical center. Health Specimens were remnant laboratory samples following routine clinical care and obtained via a Not Human Subjects Research (NHSR) classification. Therefore, informed consent was not required. Experimental Tissue Resource under HS#2012-8716 approved IRB, acted as honest broker and performed cohort identification and sample collection. Plasma drawn 1+/−days around a confirmed positive blood culture were de-identified. Clinical data such as demographics, pathology reports, hospital problem list, and antibiotic treatment at the time of blood drawn were collected and de-identified. Additional clinical parameters such as temperature, heart rate, laboratory blood test values were requested for UCI Data honest broker HS# 2012-8757, de-identified. Both healthy and patient plasma samples were aliquoted into 50 μL portions and stored at −80°C until analysis. Before use, samples were thawed at room temperature for 5 minutes. The demographics of the healthy and septic samples used in this study are summarized in Table 1.

## Author contributions

M.J. and A.J.Q. designed research; M.J. and U.U. performed research; A.S. and S.L.C. contributed assay methodology and resources; M.J., U.U., M.S., and A.J.Q. analyzed data; D.F.T. and M.S. contributed to clinical data curation and interpretation; M.J. and A.J.Q. wrote the paper; and all authors reviewed and edited the manuscript.

## Supplementary Materials

Figs. S1 to S5

Tables S1 to S11

