## Supplemental Information for "PFLISA: A New Ultrasensitive, Multiplex Immunoassay Platform Enables Cytokine Network Profiling in Sepsis"

Minsu Jang et al.

**This PDF file includes:**

Figs. S1 to S5
Tables S1 to S11

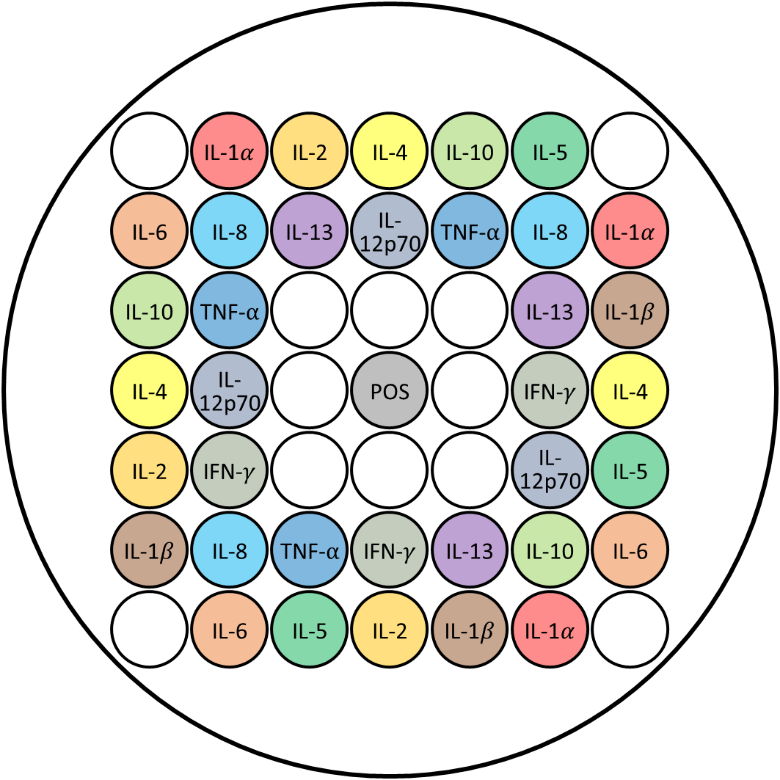

**Fig. S1.** Spatial arrangement of antibody spots corresponding to each cytokine within a single well of the 96-well plate. POS refers to the positive control spot included within each well.

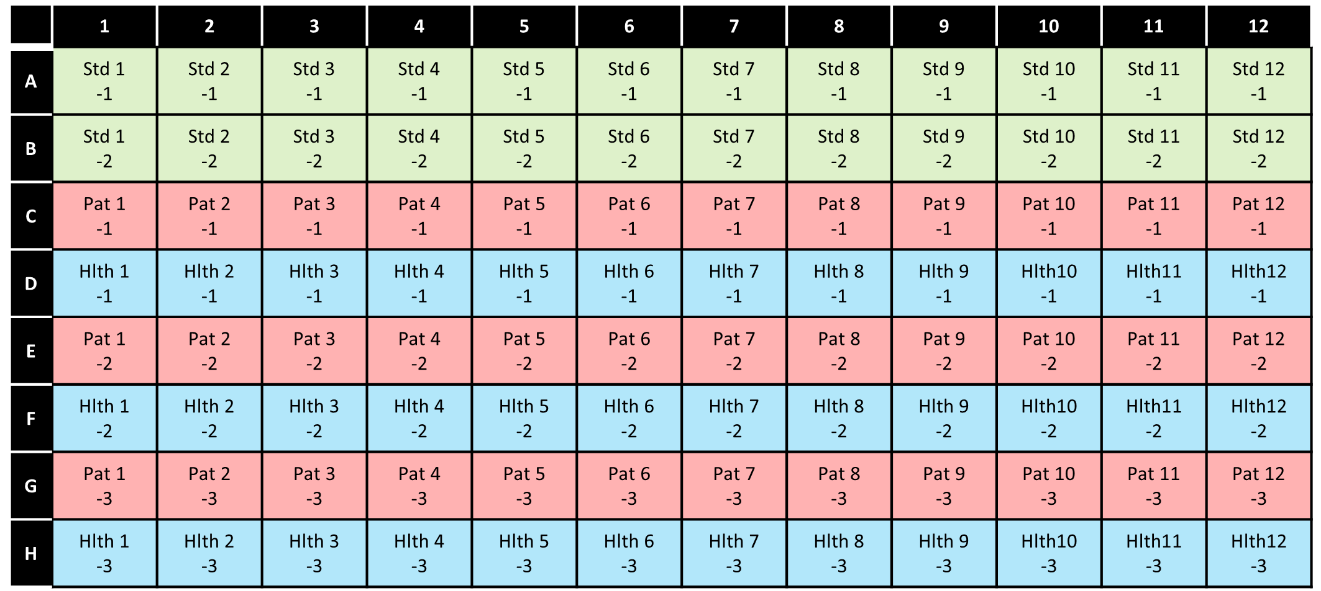

**Fig. S2.** Sample loading map of the 96-well plate for standard curves and plasma samples. (Std: Standard / Pat: Patient(Sepsis) Sample / Hlth: Healthy Sample).

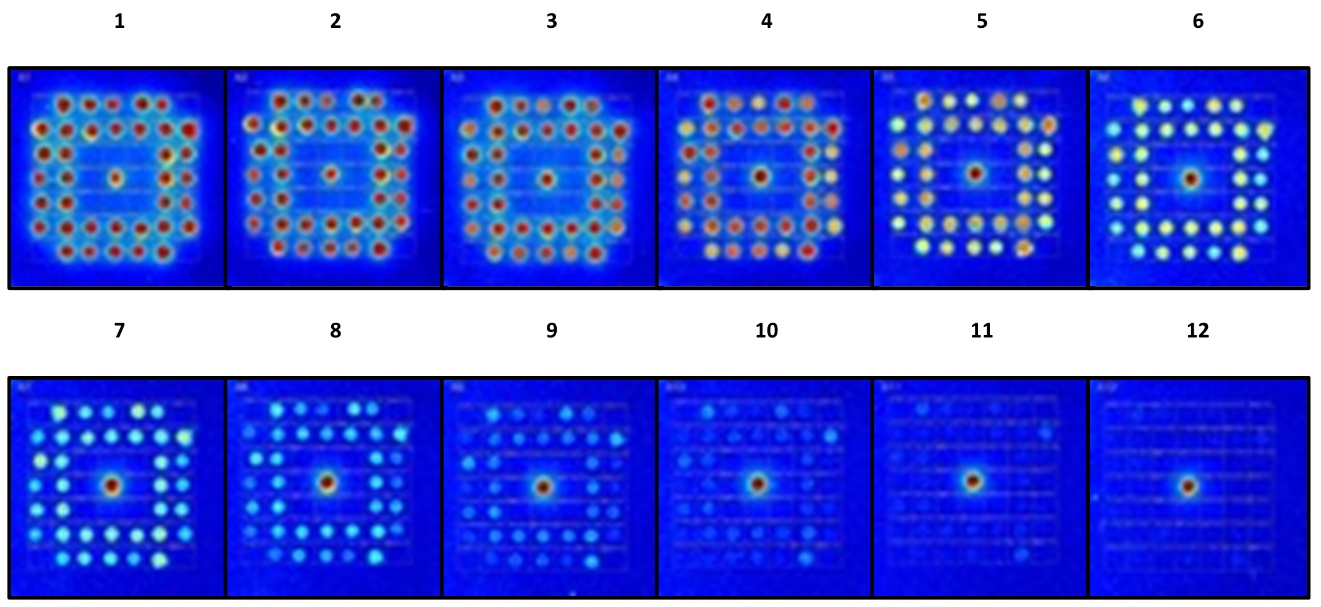

**Fig. S3.** Representative plasmonic fluorescence images of wells loaded with serially diluted standard samples.

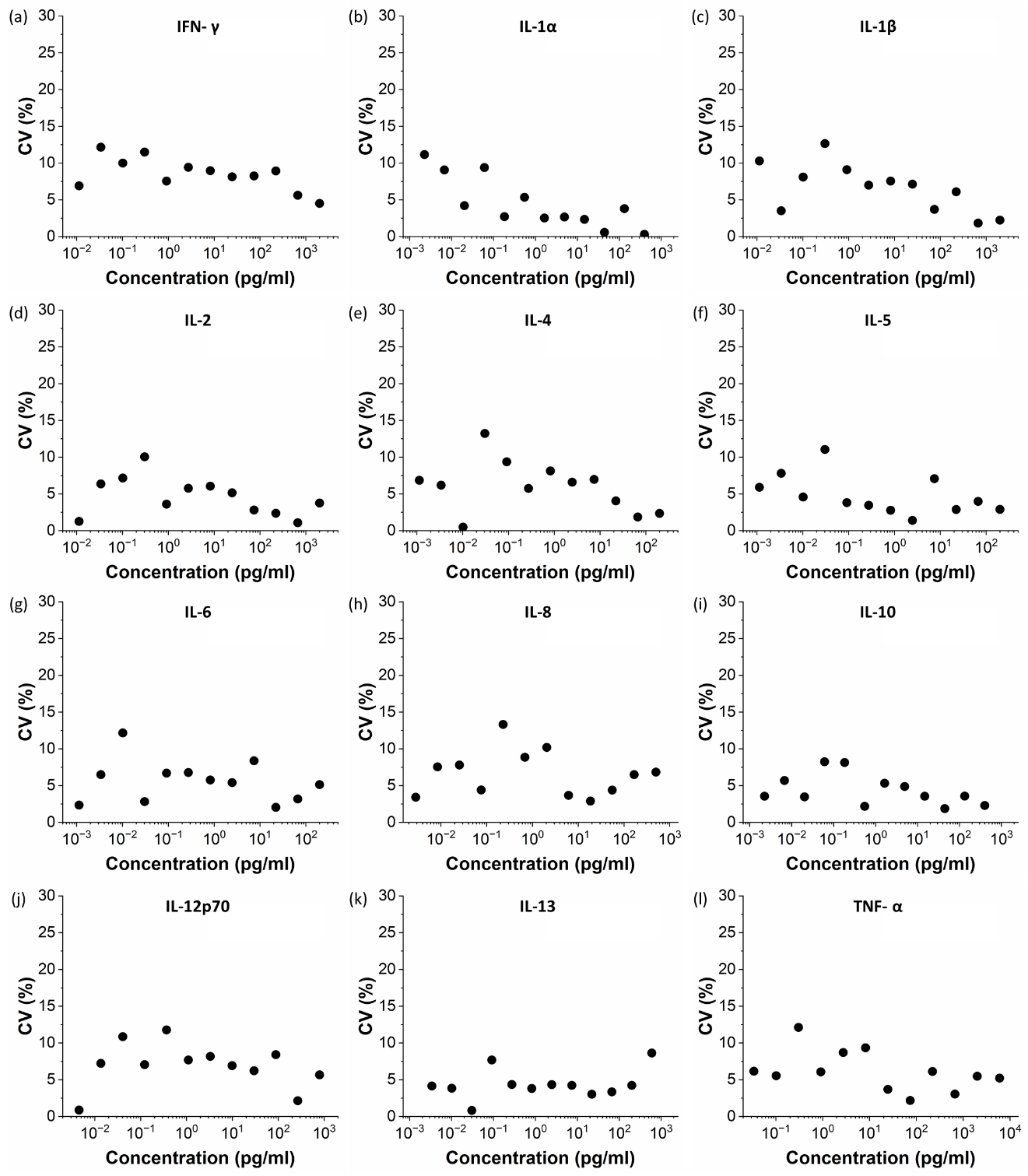

**Fig. S4.** Coefficients of variation (CVs) across three replicate spots per cytokine within a single well, demonstrating measurement reproducibility.

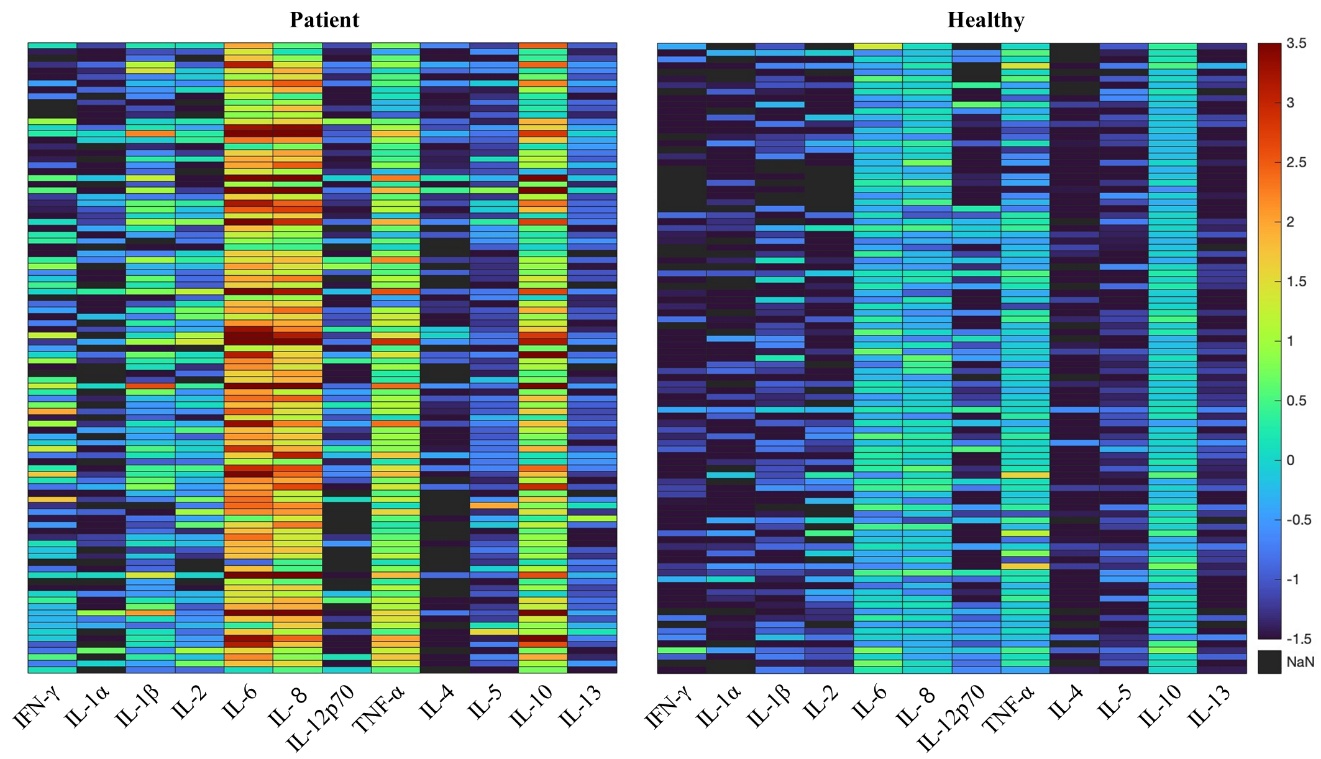

**Fig. S5.** Heatmap of cytokine concentrations for all sepsis patients (n = 100) and healthy controls (n = 97). Each row represents an individual subject, and each column represents one of the 12 target cytokines. Color scale ranges from low (blue) to high (red) concentration; black cells indicate values below the detection limit. Cytokines are ordered as pro-inflammatory (IFN-γ, IL-1α, IL-1β, IL-2, IL-6, IL-8, IL-12p70, TNF-α) followed by anti-inflammatory (IL-4, IL-5, IL-10, IL-13).

**Table S1.** Concentration of Standard 1 for individual cytokines.

| **Cytokine** | **Standard 1 Concentration (pg/mL)** |
| --- | --- |
| IFN-$\gamma$ | 600 |
| IL-1$\alpha$ | 150 |
| IL-1$\beta$ | 600 |
| IL-2 | 600 |
| IL-4 | 200 |
| IL-5 | 200 |
| IL-6 | 150 |
| IL-8 | 250 |
| IL-10 | 400 |
| IL-12p70 | 800 |
| IL-13 | 600 |
| TNF-$\alpha$ | 2,000 |

**Table S2.** Comparison of lower limits of quantification (LLOQs) between conventional cytokine detection methods and the Brightest Bio Human Inflammation 12-plex assessed in this study.

| Cytokine | LLOQ (pg/ml) | | | | |
| --- | --- | --- | --- | --- | --- |
|  | CBA | Luminex | Mesoscale | SIMOA | This study |
| IFN-𝛾 | - | 2.44 | 0.7 | 0.0391 | 0.0339 |
| IL-1α | - | - | 3.5 | 0.011 | 0.00677 |
| IL-1𝛽 | 2.4 | 1.95 | 0.56 | 0.028 | 0.0339 |
| IL-2 | - | 2.41 | 1.4 | 0.0075 | 0.0374 |
| IL-4 | - | 8.6 | 0.21 | 0.83 | 0.00339 |
| IL-5 | - | 1.95 | 0.81 | 0.0068 | 0.00339 |
| IL-6 | 1.2 | 0.73 | 0.7 | 0.0118 | 0.00339 |
| IL-8 | 4.0 | 1.22 | 0.2 | 0.608 | 0.00848 |
| IL-10 | 0.8 | 5.86 | 0.37 | 0.0068 | 0.00677 |
| IL-12p70 | 2.8 | 1.95 | 0.54 | 0.0658 | 0.0135 |
| IL-13 | - | 2.19 | 0.49 | - | 0.0102 |
| TNF-α | 1.4 | 1.71 | 1.0 | 0.051 | 0.110 |

**Table S3.** Comparison of PFLISA 12-plex standards with corresponding NIBSC/WHO cytokine reference standards. Ratios are dimensionless.

| **Cytokine** | **NIBSC/WHO Code** | **NIBSC/WHO:PFLISA Concentration Ratio** |
| --- | --- | --- |
| IL-1α | 86/636 | 1.1 |
| IL-1β | 86/552 | 1.2 |
| IL-2 | 86/500 | 0.8 |
| IL-4 | 88/656 | 1.1 |
| IL-5 | 90/586 | 1.0 |
| IL-6 | 21/308 | 1.1 |
| IL-8 | 89/520 | 1.0 |
| IL-10 | 93/722 | 1.1 |
| IL-12p70 | 95/544 | 1.0 |
| IL-13 | 94/622 | 1.0 |
| IFN-γ | 87/586 | 0.9 |
| TNF-α | 12/154 | 1.0 |

**Table S4.** Intra- and interplate precision across 16 plates.

| Cytokine | QC level | Concentration (pg/mL) | Intraplate CV (%) | Interplate CV (%) |
| --- | --- | --- | --- | --- |
| IL-1α | Low QC | 0.02 | 35.4 | 36.8 |
| IL-1α | Med QC | 0.56 | 6.9 | 11.7 |
| IL-1α | High QC | 16.23 | 5.6 | 10.6 |
| IL-1β | Low QC | 0.07 | 10.7 | 15.4 |
| IL-1β | Med QC | 2.07 | 6.5 | 12.9 |
| IL-1β | High QC | 59.94 | 6.5 | 9.4 |
| IL-2 | Low QC | 0.08 | 13.0 | 14.8 |
| IL-2 | Med QC | 2.02 | 4.8 | 10.1 |
| IL-2 | High QC | 62.19 | 6.3 | 13.8 |
| IL-4 | Low QC | 0.03 | 8.8 | 12.2 |
| IL-4 | Med QC | 0.68 | 4.5 | 8.9 |
| IL-4 | High QC | 19.66 | 3.4 | 9.6 |
| IL-5 | Low QC | 0.03 | 10.4 | 15.7 |
| IL-5 | Med QC | 0.67 | 5.0 | 10.0 |
| IL-5 | High QC | 19.22 | 4.9 | 11.7 |
| IL-6 | Low QC | 0.02 | 5.5 | 11.1 |
| IL-6 | Med QC | 0.53 | 4.4 | 9.4 |
| IL-6 | High QC | 15.55 | 4.3 | 11.6 |
| IL-8 | Low QC | 0.03 | 8.4 | 9.1 |
| IL-8 | Med QC | 0.82 | 8.6 | 10.7 |
| IL-8 | High QC | 24.56 | 3.4 | 12.2 |
| IL-10 | Low QC | 0.05 | 6.7 | 11.3 |
| IL-10 | Med QC | 1.44 | 3.8 | 11.1 |
| IL-10 | High QC | 41.46 | 5.9 | 14.5 |
| IL-12p70 | Low QC | 0.10 | 9.9 | 12.1 |
| IL-12p70 | Med QC | 2.66 | 4.5 | 12.2 |
| IL-12p70 | High QC | 71.23 | 4.5 | 12.7 |
| IL-13 | Low QC | 0.02 | 14.6 | 11.6 |
| IL-13 | Med QC | 0.63 | 6.9 | 13.1 |
| IL-13 | High QC | 18.81 | 4.0 | 11.1 |
| IFN-γ | Low QC | 0.09 | 9.7 | 18.9 |
| IFN-γ | Med QC | 1.98 | 4.7 | 9.0 |
| IFN-γ | High QC | 63.39 | 5.2 | 10.2 |
| TNF-α | Low QC | 0.21 | 11.4 | 18.2 |
| TNF-α | Med QC | 5.68 | 7.4 | 15.0 |
| TNF-α | High QC | 185.26 | 5.9 | 12.0 |

**Table S5.** Matrix spike recovery at low, medium, and high concentrations for plasma.

| Analyte | Level | Spike Conc. (pg/mL) | Avg. % Recovery (n = 4) | % Recovery Range |
| --- | --- | --- | --- | --- |
| IL-1α | Low | 0.38 | 58.5 | 23.3—90.2 |
|  | Medium | 3.75 | 61.8 | 24.6—89.0 |
|  | High | 18.75 | 61.7 | 20.6—90.4 |
| IL-1β | Low | 1.50 | 78.3 | 54.1—87.3 |
|  | Medium | 15.00 | 83.2 | 71.1—92.9 |
|  | High | 75.00 | 76.5 | 63.9—82.3 |
| IL-2 | Low | 1.50 | 62.9 | 51.9—76.8 |
|  | Medium | 15.00 | 70.4 | 57.1—85.7 |
|  | High | 75.00 | 74.8 | 56.4—90.2 |
| IL-4 | Low | 0.50 | 79.3 | 68.8—86.1 |
|  | Medium | 5.00 | 79.3 | 69.9—85.7 |
|  | High | 25.00 | 84.4 | 69.8—96.2 |
| IL-5 | Low | 0.50 | 80.7 | 71.4—91.1 |
|  | Medium | 5.00 | 85.1 | 82.0—88.6 |
|  | High | 25.00 | 90.8 | 80.1—98.6 |
| IL-6 | Low | 0.38 | 77.5 | 62.6—89.5 |
|  | Medium | 3.75 | 84.8 | 72.5—91.6 |
|  | High | 18.75 | 84.3 | 67.0—99.9 |
| IL-8 | Low | 0.63 | 82.1 | 39.4—136.8 |
|  | Medium | 6.25 | 89.2 | 74.9—102.1 |
|  | High | 31.25 | 112.1 | 102.1—119.0 |
| IL-10 | Low | 1.00 | 67.5 | 49.0—82.6 |
|  | Medium | 10.00 | 68.8 | 51.0—85.2 |
|  | High | 50.00 | 66.6 | 44.0—84.4 |
| IL-12p70 | Low | 2.00 | 75.8 | 61.3—90.9 |
|  | Medium | 20.00 | 79 | 59.9—90.5 |
|  | High | 100.00 | 89.9 | 51.3—115.3 |
| IL-13 | Low | 0.50 | 74.1 | 63.4—83.0 |
|  | Medium | 5.00 | 79.9 | 72.3—88.5 |
|  | High | 25.00 | 76.6 | 62.8—89.1 |
| IFN-γ | Low | 1.50 | 76.4 | 64.0—83.5 |
|  | Medium | 15.00 | 79.6 | 72.3—88.5 |
|  | High | 75.00 | 78.9 | 70.8—83.9 |
| TNF-α | Low | 5.00 | 72.2 | 63.0—84.0 |
|  | Medium | 50.00 | 84.5 | 76.0—90.5 |
|  | High | 250.00 | 80.3 | 62.9—96.6 |

**Table S6.** Matrix spike recovery at low, medium, and high concentrations for serum.

| Analyte | Level | Spike Conc. (pg/mL) | Avg. % Recovery (n = 4) | % Recovery Range |
| --- | --- | --- | --- | --- |
| IL-1α | Low | 0.38 | 70.3 | 49.5—89.8 |
|  | Medium | 3.75 | 67.1 | 39.5—85.4 |
|  | High | 18.75 | 72.2 | 48.8—93.4 |
| IL-1β | Low | 1.50 | 72.4 | 47.6—83.8 |
|  | Medium | 15.00 | 76.8 | 56.6—88.6 |
|  | High | 75.00 | 71.4 | 50.8—86.4 |
| IL-2 | Low | 1.50 | 60.4 | 47.0—72.8 |
|  | Medium | 15.00 | 64.1 | 57.6—73.6 |
|  | High | 75.00 | 67.9 | 60.5—76.8 |
| IL-4 | Low | 0.50 | 74.5 | 55.5—85.7 |
|  | Medium | 5.00 | 72.5 | 51.2—83.9 |
|  | High | 25.00 | 77 | 51.9—91.2 |
| IL-5 | Low | 0.50 | 69.8 | 37.2—89.7 |
|  | Medium | 5.00 | 71.6 | 40.4—86.4 |
|  | High | 25.00 | 74.7 | 41.8—89.1 |
| IL-6 | Low | 0.38 | 83.4 | 70.3—95.4 |
|  | Medium | 3.75 | 83.2 | 73.6—89.4 |
|  | High | 18.75 | 81.3 | 67.1—92.2 |
| IL-8 | Low | 0.63 | 72.9 | 41.1—100.1 |
|  | Medium | 6.25 | 72.6 | 55.8—99.2 |
|  | High | 31.25 | 94.9 | 76.1—113.6 |
| IL-10 | Low | 1.00 | 65.5 | 44.4—83.7 |
|  | Medium | 10.00 | 65.8 | 54.8—84.2 |
|  | High | 50.00 | 64.4 | 53.3—90.2 |
| IL-12p70 | Low | 2.00 | 81.8 | 67.1—93.0 |
|  | Medium | 20.00 | 81 | 72.6—89.8 |
|  | High | 100.00 | 93.3 | 85.5—98.0 |
| IL-13 | Low | 0.50 | 57.1 | 38.7—74.8 |
|  | Medium | 5.00 | 56.5 | 38.6—77.04 |
|  | High | 25.00 | 59.6 | 42.5—77.7 |
| IFN-γ | Low | 1.50 | 80 | 65.4—90.2 |
|  | Medium | 15.00 | 81.5 | 68.5—90.1 |
|  | High | 75.00 | 81.6 | 74.0—94.8 |
| TNF-α | Low | 5.00 | 69.6 | 54.4—84.5 |
|  | Medium | 50.00 | 75.2 | 66.2—85.8 |
|  | High | 250.00 | 79.8 | 66.6—93.6 |

**Table S7.** Individual demographic, clinical, and physiological characteristics of the enrolled patients with sepsis (n = 100).

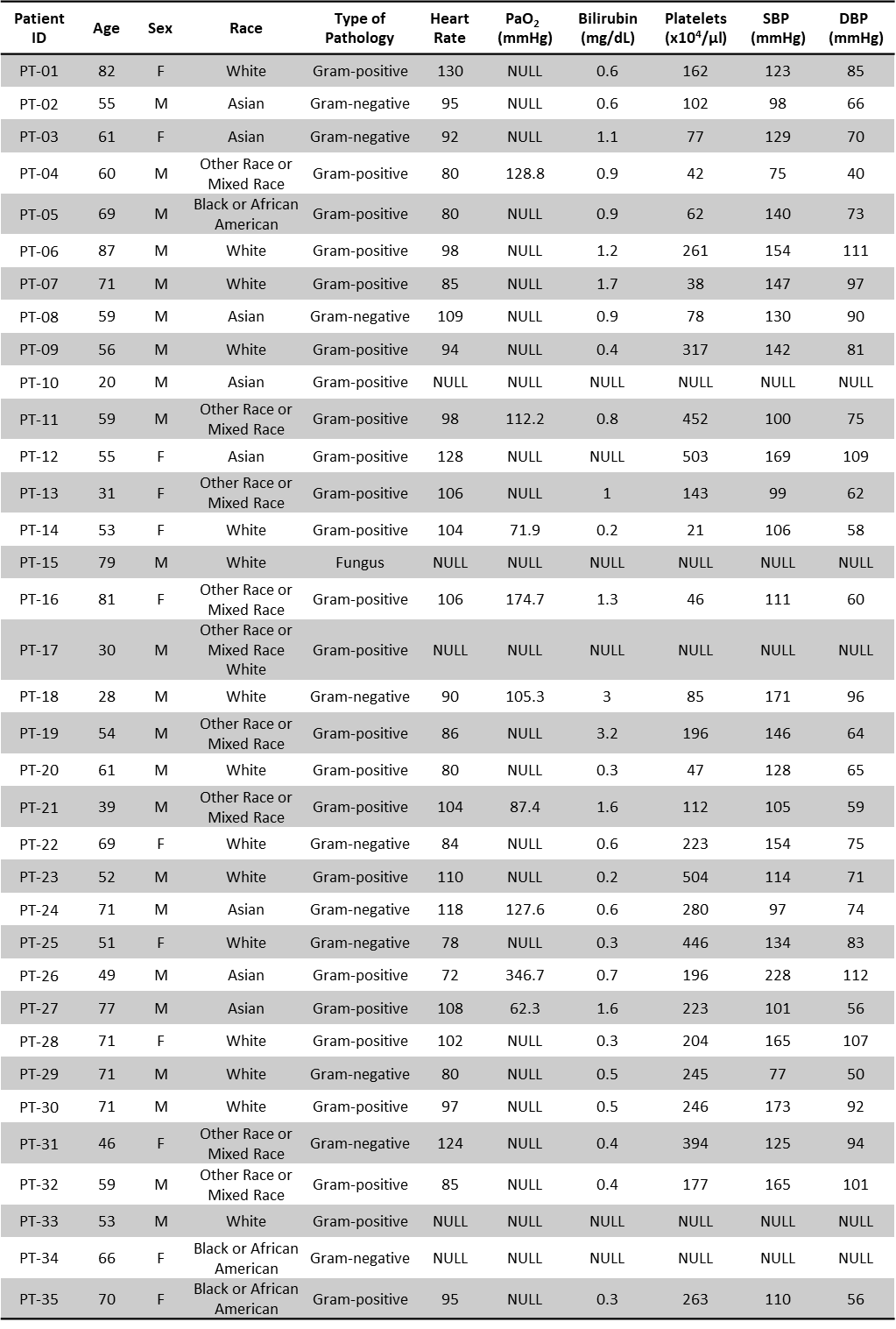

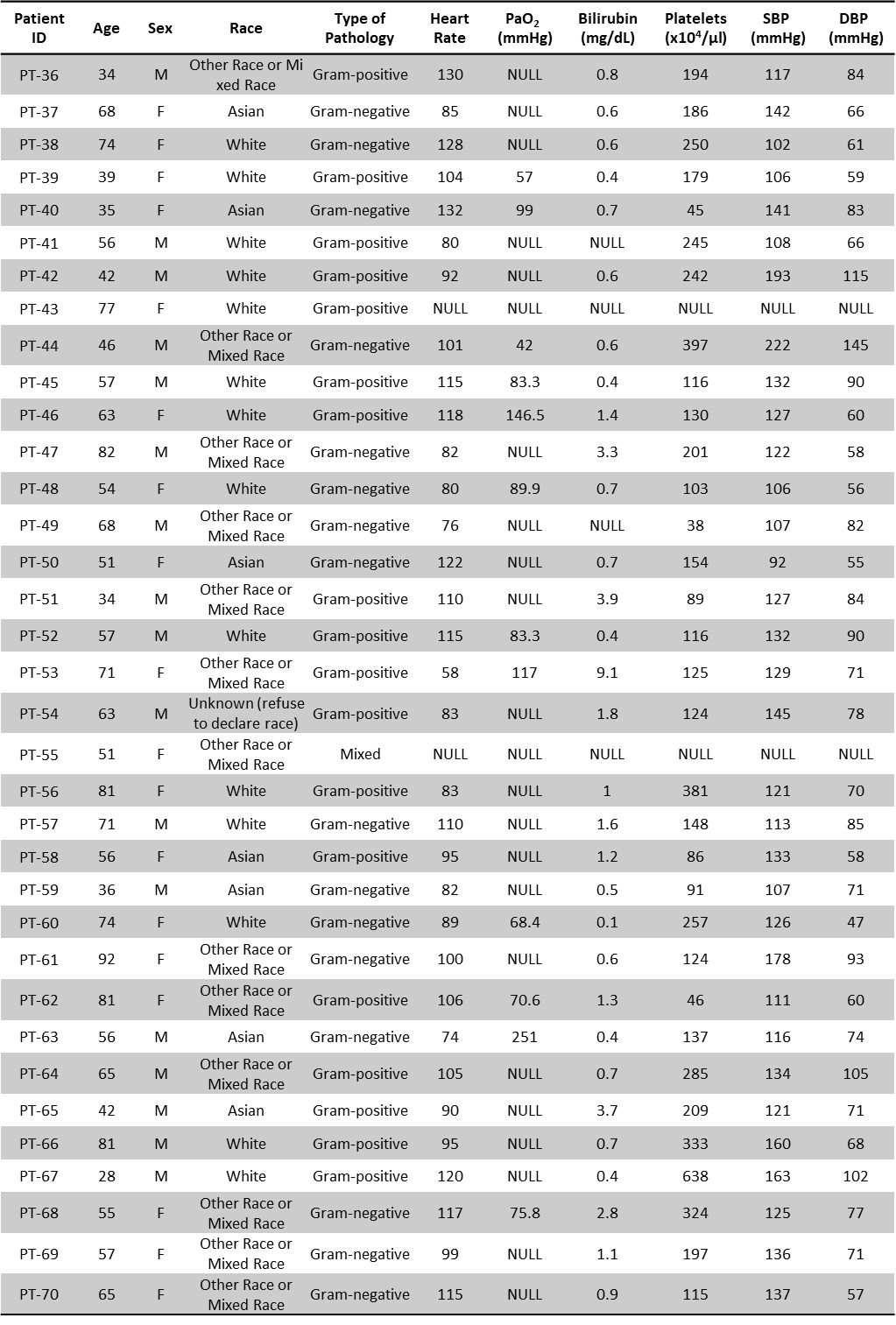

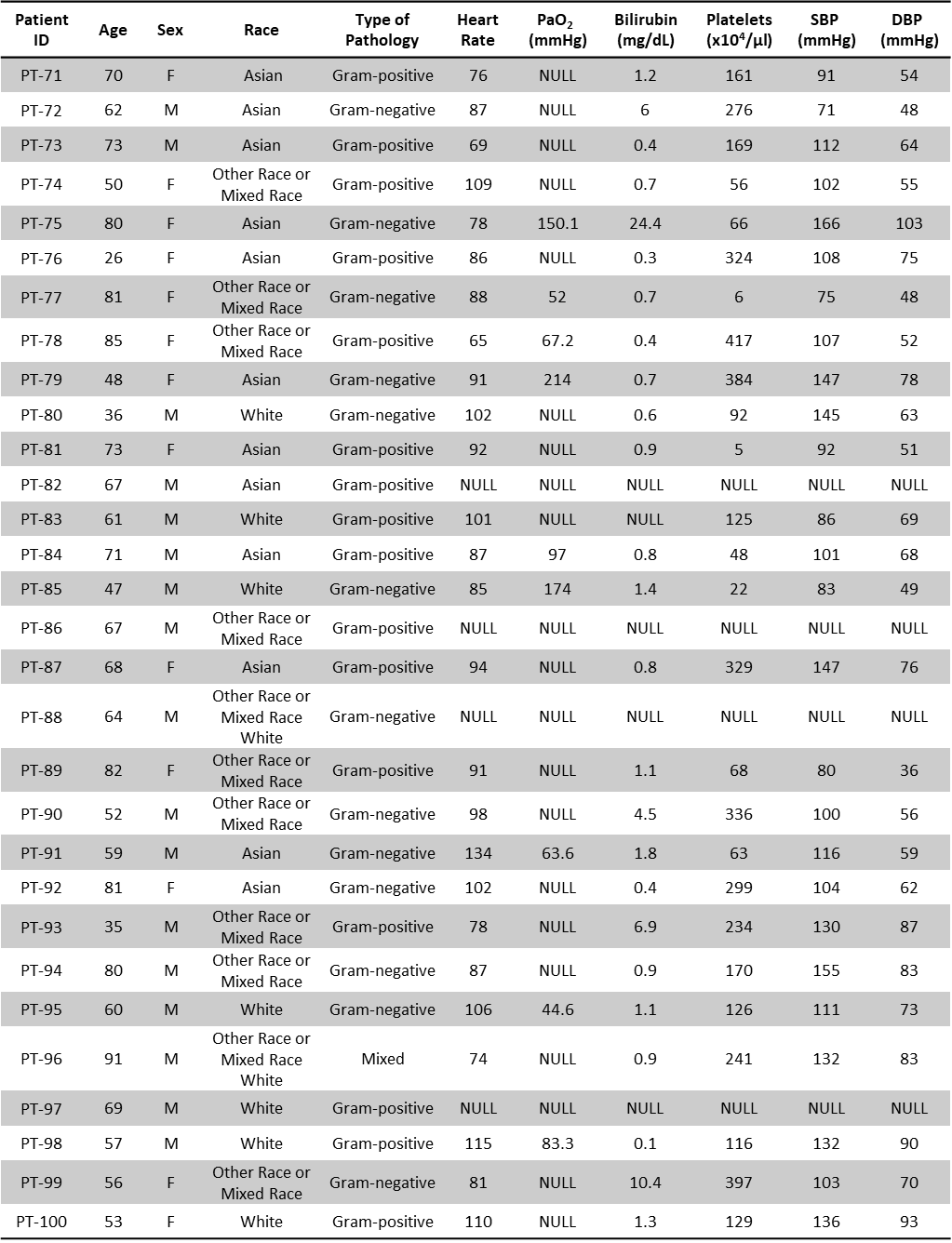

**Table S8.** Cytokine concentrations measured by PFLISA in sepsis patients and healthy controls.

|  | **Sepsis [IQR]** | **Healthy [IQR]** |
| --- | --- | --- |
| IFN-γ | 0.341 [0.039-1.61] | 0.029 [0.012-0.066] |
| IL-1α | 0.070 [0.022-0.318] | 0.026 [0.009-0.079] |
| IL-1β | 0.598 [0.177-2.00] | 0.070 [0.029-0.166] |
| IL-2 | 0.456 [0.138-1.28] | 0.053 [0.021-0.175] |
| IL-4 | 0.044 [0.023-0.130] | 0.019 [0.010-0.036] |
| IL-5 | 0.100 [0.055-0.216] | 0.049 [0.023-0.088] |
| IL-6 | 62.1 [24.8-232.2] | 0.791 [0.408-1.51] |
| IL-8 | 43.9 [15.7-171.8] | 0.839 [0.463-1.37] |
| IL-10 | 10.1 [4.03-60.6] | 0.928 [0.705-1.36] |
| IL-12p70 | 0.078 [0.028-0.278] | 0.071 [0.025-0.319] |
| IL-13 | 0.113 [0.059-0.214] | 0.043 [0.020-0.063] |
| TNF-α | 6.57 [2.94-18.6] | 0.947 [0.461-1.53] |

**Table S9.** Cytokine concentrations measured by PFLISA in pathogen type (gram positive and negative).

|  | **Gram Positive [IQR]** | **Gram Negative [IQR]** | **FDR p-value** |
| --- | --- | --- | --- |
| IFN-γ | 0.164 [0.032-1.69] | 0.600 [0.093-1.29] | 0.273 |
| IL-1α | 0.063 [0.02-0.189] | 0.071 [0.023-0.339] | 0.493 |
| IL-1β | 0.436 [0.138-1.708] | 0.631 [0.259-4.32] | 0.225 |
| IL-2 | 0.392 [0.111-1.099] | 0.482 [0.145-1.24] | 0.493 |
| IL-4 | 0.040 [0.024-0.115] | 0.047 [0.022-0.135] | 0.751 |
| IL-5 | 0.097 [0.055-0.255] | 0.100 [0.055-0.207] | 0.968 |
| IL-6 | 56.32 [15.9-111.4] | 120.5 [41.2-2171.1] | 0.022 |
| IL-8 | 27.7 [9.21-79.2] | 82.3 [22.8-288.2] | 0.022 |
| IL-10 | 7.48 [2.74-27.3] | 14.4 [6.80-339.0] | 0.022 |
| IL-12p70 | 0.069 [0.018-0.533] | 0.089 [0.039-0.221] | 0.968 |
| IL-13 | 0.110 [0.044-0.229] | 0.121 [0.039-0.221] | 0.493 |
| TNF-α | 4.408 [2.08-10.6] | 10.7 [3.75-37.0] | 0.022 |

**Table S10.** Cytokine concentrations measured by PFLISA in survivors and non-survivors.

|  | **Survivors [IQR]** | **Non-survivors [IQR]** | **p-value** |
| --- | --- | --- | --- |
| IFN-γ | 0.761 [0.056-2.19] | 0.192 [0.031-0.607] | 0.030 |
| IL-1α | 0.064 [0.023-0.203] | 0.090 [0.022-0.346] | 0.732 |
| IL-1β | 0.656 [0.177-1.85] | 0.579 [0.222-2.50] | 0.913 |
| IL-2 | 0.520 [0.135-1.31] | 0.411 [0.154-1.18] | 0.752 |
| IL-4 | 0.047 [0.024-0.125] | 0.042 [0.012-0.138] | 0.839 |
| IL-5 | 0.097 [0.055-0.217] | 0.104 [0.056-0.213] | 0.775 |
| IL-6 | 49.9 [20.8-148.5] | 131.2 [54.9-292.2] | 0.011 |
| IL-8 | 28.6 [10.3-91.8] | 82.5 [25.0-189.7] | 0.011 |
| IL-10 | 9.26 [2.65-60.9] | 14.4 [5.29-47.9] | 0.288 |
| IL-12p70 | 0.095 [0.035-0.362] | 0.052 [0.018-0.162] | 0.165 |
| IL-13 | 0.113 [0.080-0.201] | 0.121 [0.046-0.292] | 0.908 |
| TNF-α | 6.21 [3.02-19.1] | 7.69 [2.85-13.5] | 0.936 |

**Table S11.** Cytokine concentrations measured by PFLISA in organ dysfunction severity.

|  | **No Dysfunction [IQR]** | **Single-organ Dysfunction [IQR]** | **Multi-organ Dysfunction [IQR]** | **p-value** |
| --- | --- | --- | --- | --- |
| IFN-γ | 0.237 [0.040-2.13] | 0.594 [0.054-1.50] | 0.249 [0.024-1.18] | 0.464 |
| IL-1α | 0.064 [0.011-0.246] | 0.070 [0.026-0.149] | 0.079 [0.026-0.346] | 0.716 |
| IL-1β | 0.730 [0.192-1.70] | 1.27 [0.288-3.09] | 0.554 [0.148-1.84] | 0.351 |
| IL-2 | 0.174 [0.066-0.583] | 0.698 [0.335-1.81] | 0.523 [0.203-1.37] | 0.011 |
| IL-4 | 0.040 [0.014-0.088] | 0.039 [0.021-0.123] | 0.126 [0.043-0.159] | 0.135 |
| IL-5 | 0.066 [0.055-0.285] | 0.153 [0.069-0.238] | 0.100 [0.056-0.172] | 0.585 |
| IL-6 | 50.2 [23.0-109.8] | 125.1 [55.9-1115.9] | 113.3 [40.7-223.5] | 0.069 |
| IL-8 | 22.1 [13.6-61.7] | 79.4 [31.3-223.1] | 103.8 [33.7-273.4] | 0.010 |
| IL-10 | 7.89 [2.61-21.7] | 14.2 [8.57-84.0] | 21.0 [6.22-61.6] | 0.036 |
| IL-12p70 | 0.078 [0.042-0.310] | 0.095 [0.037-0.311] | 0.106 [0.020-0.722] | 0.962 |
| IL-13 | 0.090 [0.044-0.192] | 0.134 [0.094-0.206] | 0.134 [0.057-0.289] | 0.284 |
| TNF-α | 4.34 [2.46-14.0] | 10.5 [4.94-27.9] | 6.22 [2.93-13.3] | 0.167 |
